# Prior heat stress increases pathogen susceptibility in the model cnidarian *Exaptasia diaphana*

**DOI:** 10.1101/2023.10.12.562090

**Authors:** Sofia C. Diaz de Villegas, Erin M. Borbee, Peyton Y. Abdelbaki, Lauren E. Fuess

## Abstract

Anthropogenic climate change has significantly altered terrestrial and marine ecosystems globally, often in the form of climate-related events such as thermal anomalies and disease outbreaks. Although the isolated effects of these stressors have been well documented, a growing body of literature suggests that stressors often interact, resulting in complex effects on ecosystems, including coral reefs where sequential associations between heat stress and disease have had profound impacts. Here we used the model cnidarian *Exaiptasia diaphana* to investigate mechanisms linking prior heat stress to increased disease susceptibility. We examined anemone pathogen susceptibility and physiology (symbiosis, immunity, and energetics) following recovery from heat stress. We observed significantly increased pathogen susceptibility in anemones previously exposed to heat stress. Notably, prior heat stress reduced anemone energetic reserves (carbohydrate concentration), and activity of multiple immune components. Minimal effects of prior heat stress on symbiont density were observed. Together, results suggest changes in energetic availability might have the strongest effect on pathogen susceptibility and immunity following heat stress. The results presented here provide critical insight regarding the interplay between heat stress recovery and pathogen susceptibility in cnidarians and are an important first step towards understanding temporal associations between these stressors.

## Introduction

Anthropogenic pressure has led to environmental and ecological changes in terrestrial and aquatic ecosystems across the world, often in the form of climate-related events.^1,2^ Marine ecosystems, in particular, have been significantly affected by multiple climate-induced stressors, such as ocean warming and disease outbreaks^3–5^. Both of these stressors directly affect organismal physiology through alteration of metabolic function, and indirectly affect a variety of ecological processes, such as oxygen availability, nutrient cycling, and ocean circulation^3,6^. As such, ocean warming, specifically marine heatwaves, are a leading cause of mass mortality in marine organisms^7–10^. Similarly, increases in the frequency and severity of disease outbreaks pose significant threats to marine ecosystems. Marine taxa are affected by a large diversity of pathogens ranging from viruses and bacteria to fungi and metazoans^11^, many of which have triggered significant mortality events in a variety of marine organisms^12–15^.

Although the isolated effects of ocean warming, disease outbreaks, and other climate-associated stressors have been well documented within marine ecosystems^3,5^, a growing body of literature suggests that these stressors often coincide. Depending on the circumstances and stressors in question, interactions may range from cumulative to synergistic, or even antagonistic in nature.^16,17^ Associations between heat stress and disease outbreaks are particularly prevalent and have often contributed to extensive marine mortality.^18^ Consequently, studies have attempted to improve understanding of associations between heat stress and disease in marine ecosystems, but most have focused only on simultaneous synergy between the two stressors.^19–21^ However, frequent increases in disease outbreaks following marine heatwaves suggest that these two stressors may more commonly accumulate in a sequential, rather than simultaneous, manner.^22–25^ Despite these observations, the mechanisms driving sequential temporal patterns between heat stress and disease are poorly understood and necessitate further investigation.

Coral reef ecosystems present a unique opportunity to investigate sequential associations between heat stress and disease. These ecosystems are commonly faced with a variety of climate stressors^26^, which have resulted in unprecedented declines in coral cover over the past several decades.^27,28^ Increasing sea surface temperature (SST), in particular, has exacerbated the frequency and severity of coral decline. Two major drivers of these declines are heat-induced bleaching—loss of endosymbiotic algae^29,30^—and disease^31^. Although isolated heat-induced bleaching^32–34^ and disease are common^35–38^, these events have also been frequently observed in close spatial and temporal associations.^39–43^ In many cases, coral disease follows bleaching^25^, with extensive disease-related mortality often reported in the weeks to months after heat stress.^24,44^ Despite these observations, nearly all experimental studies have focused on the synergistic effects of simultaneous heat stress and disease, specifically documenting increases in disease susceptibility when these stressors occur at the same time.^45,46^ Synchronous associations between heat stress and increased pathogen susceptibility may be linked to heat induced suppression of immunity,^47–51^ though other studies have instead demonstrated minimal or positive effects of heat stress on coral immunity.^52–54^ Furthermore, it is unclear if similar mechanisms underly sequential associations between heat stress and disease outbreaks.

Sequential associations between heat stress and disease outbreaks may be mediated by several mechanisms. First, heat stress may induce changes in host energetic reserves, which lead to immunosuppression. The loss of symbionts which typically occurs as a result of heat stress may severely limit the energetic resources available to the host.^55–57^ According to resource allocation theory, organisms must distribute a finite amount of resources amongst diverse biological processes—growth, reproduction, metabolism, immunity, etc.^58–60^ This may involve trade-offs where resources are shifted between competing needs in order to maximize fitness.^58–60^ Consequently, reduction in the total amount of resources may alter the proportions in which resources are distributed, potentially resulting in the suppression of one or more biological processes. As such, immune suppression is one of the most commonly documented energetic trade-offs associated with resource limitation.^61,62^ Heat-stress induced limitations in energetics may exacerbate resource tradeoffs and result in changes to resource allocation, including allocation to host immune responses. Alternatively, immune-energetic trade-offs may also occur in the absence of resource limitation in response to biotic or abiotic pressures: organisms may dynamically shift resources between processes in response to pressing needs caused by environmental stress or other changes in biological condition.^63^

Alternatively, dynamic changes in symbiosis during heat stress and associated recovery may contribute to observed links between heat stress and disease outbreaks. Suppression of host immunity by symbionts has been hypothesized as an important mechanism required for the successful establishment and maintenance of symbiotic relationships.^64^ Previous laboratory studies have confirmed negative associations between symbiosis and immunity across several cnidarians.^65–68^ However, the dynamics and consequences of immune-symbiosis interplay during the breakdown and re-establishment of symbiosis remain unclear. One possibility that the rapid repopulation of endosymbionts following heat-induced dysbiosis exacerbates symbiotic immunosuppression, potentially explaining, in part, naturally observed patterns of disease after heat-induced bleaching events.

Here we describe the results of a study aimed at assessing the sequential associations between heat stress and disease outbreaks in cnidarians, with a specific focus on the roles of changes in energetics, symbiont density, and host immunity in these processes. In order to comprehensively address this question, we leveraged the powerful emergent cnidarian model species, *Exaiptasia diaphana* (common name Aiptasia). While tropical scleractinian corals are exceptionally important from an ecological standpoint, their precarious species statuses and inherent experimental limitations complicate the laboratory investigation of complex biological phenomenon in these species. In contrast with tropical corals, Aiptasia is easily reared, maintained, and destructively sampled in large amounts without many of the experimental and ethical challenges of working with tropical corals.^69^ Aiptasia also maintains symbiotic relationships with endosymbiotic algae of the family Symbiodiniaceae, like tropical corals.^70,71^ Finally, Aiptasia is a particularly handy system due to the existence of established, genetically identical clonal lines, allowing for control of genetic variation in experimentation.^69^ We leveraged the *E. diaphana* model system to investigate the mechanisms linking heat stress and disease susceptibility in cnidarians. Specifically, we examined host immunity, symbiont density, and energetic reserves following heat-stress, as well as the effects of prior heat stress on pathogen susceptibility, in two genetically distinct clonal lines of *E. diaphana*. We document a significant association between prior heat stress and subsequent increased pathogen susceptibility and highlight the potential physiological mechanisms of this association. The results described are an important first step towards understanding temporal links between two of the most significant climate-associated stressors faced by cnidarians.

## Methods

### Experimental Design

Anemones were obtained from laboratory-maintained clonal lines of symbiotic *E. diaphana* (VWB9 and H2; V. Weis). Anemones from each clonal line were randomly assigned to one of four possible treatment combinations: ambient temperature and placebo, heat stress and placebo, ambient temperature and infection, or heat stress and infection (Table 1). Individuals were transferred accordingly into sterile, lidded 6-well culture plates filled with 10 mL of filter-sterile artificial sea water (FASW). Each well contained a single anemone and plates were distributed such that each treatment combination had 30 plates. Plates assigned to ambient temperature treatment were maintained in a single temperature-controlled incubator (Percival Scientific; Model AL-41L4) at 27°C for the duration of the experiment. Plates assigned to heat stress treatment were randomly divided between two separate temperature-controlled incubators (VWR; Model 2005, VWR; Model 3734). Prior to the start of the experiment, all individuals were acclimated to well plates for four days at 27°C in their respective incubators. Throughout the entire experiment, anemones in all three incubators were maintained under the same light conditions (12hr:12hr light: dark cycles with approximately 40-60 µmol·m^-2^ PAR) and received bi-weekly feeding (*Artemia nauplii)* and subsequent water changes until the point of infection. Plate locations were randomized within and between incubators at the time of water changes, as appropriate.

**Table 1:** Initial anemone distribution for H2 and VWB9 conal lines across each of four treatment combinations.

|  | VWB9 |  | H2 |  |
| --- | --- | --- | --- | --- |
|  | Ambient | Heat | Ambient | Heat |
| Placebo | 86 | 86 | 86 | 86 |
| Pathogen | 86 | 86 | 86 | 86 |

### Experimental Heat Stress & Pathogen Challenge

Heat stressed anemones were exposed to a temperature profile which consisted of a seven-day ramping period wherein temperatures were raised 1°C per day to a final temperature of 34°C which was maintained for five days before an immediate return to ambient temperatures (27°C) on day six. The chosen protocol was developed based on previous studies that used 34°C as a target temperature^45,72,73^ and pilot data which demonstrated this regime induced bleaching (measured as decline in chlorophyll fluorescence) with limited mortality. This approach was employed to mimic a sublethal bleaching event from which individuals recover so as to test our hypotheses regarding the impact of reestablishment of symbiosis on host immunity. During the experiment, bleaching was confirmed by visual assessment of anemone color and was found to be consistent with the expected timeline. Following heat stress, anemones were immediately returned to 27°C and allowed to recover for two weeks. At the conclusion of this recovery period, surviving anemones from both temperature treatments were inoculated with either a pathogen or a placebo (FASW) by plate, according to *a priori* assignments. Pathogen exposure was conducted using the known coral pathogen *V. coralliilyticus* BAA 450^74^ at a predetermined dosage of 1 x 10^8^ CFU·mL^-1^.^45^ Final bacterial concentrations were confirmed with serial dilutions and plating. Following exposure, a subset of anemones was monitored for survivorship (**Table 2)**: mortality data was collected every two hours for the first 24 hours, and then daily for the remainder of the experiment (96 hours post-infection) at which point 89% of exposed anemones had died. Individuals were considered dead when they exhibited more than 75% tissue degradation or had 75% tissue degradation for more than 24 hours.

**Table 2:** Final sample sizes for physiological and survivorship analyses.

| Survivorship | VWB9 |  | H2 |  |
| --- | --- | --- | --- | --- |
|  | Ambient | Heat | Ambient | Heat |
| Placebo | 30 | 26 | 19 | 21 |
| Infection | 30 | 25 | 20 | 13 |
| Physiology | VWB9 |  | H2 |  |
|  | Ambient | Heat | Ambient | Heat |
| Placebo | 14 | 13 | 12 | 11 |
| Pathogen | 17 | 12 | 11 | 10 |

### Sample Collection and Processing

Twelve hours after inoculation, a second subset of randomly selected anemones was sampled for the assessment of symbiont density, immune activity, and energetic reserves (**Table 2**). Anemones were anesthetized in 10 mL of MgCl_2_ (0.37 M) prior to sampling. First, we excised a single tentacle for estimates of symbiont density described below. Subsequent tentacle processing was modified from existing protocols.^75^ Following tentacle excision, the remaining anemone was halved using a sterile razor blade; one half was flash frozen for the immune and energetic assays described herein. Frozen tissue samples were homogenized with 3 mm glass beads in 1.5 mL of 100 mM Tris + 0.05 M DTT buffer (pH 7.8) for 1 minute at 5 m·s^-1^ (Tim Bateman, *pers. comm*.) using a bead mill (Fisherbrand). For each sample, a 200 µL aliquot of homogenate was transferred into a separate microcentrifuge tube and stored at −20°C for melanin assays. The remaining homogenate for each sample was then transferred into a new 2 mL microcentrifuge tube and centrifuged at 2250xg at 4°C for five minutes. Resulting supernatants were aliquoted accordingly for each set of assays (protein, lipid, carbohydrates, ash-free dry weight) and stored at −80°C (protein fraction) or −20°C (all other fractions) prior to downstream analyses.

### Symbiont Density Estimation

Mounted tentacles were imaged using brightfield and fluorescence microscopy on the Cytation1 Cell Imaging Multimode Reader (Biotek). Approximate tentacle area was determined via brightfield microscopy. Symbiont autofluorescence was measured using a chlorophyll a fluorescent filter (Agilent; EX 445/45 nm, EM 685/40 nm). Brightfield and fluorescent images of each tentacle were combined to standardize fluorescence by tentacle area. The average pixel intensity of the chlorophyll a fluorescence within each standardized tentacle area was used as a proxy for symbiont density.

### Immune Assays

Cnidarian immunity was measured using a suite of standardized immune assays originally developed for scleractinian corals^76,77^ and adapted for use in *E. diaphana*. We measured responses related to three major innate immune pathways: antioxidant, phenoloxidase, and antibacterial pathways. All immune assays were run in triplicate on separate 96-well plates and measured using the Cytation1 Cell Imaging Multimode Reader and Gen5 Software (Biotek). Negative controls for all assays were 100mM Tris + 0.05 mM DTT buffer (homogenization buffer). All enzymatic immune assays were standardized by protein concentration using established protocols for Red660 colorimetric assays^76,77^, while melanin concentration was standardized by desiccated tissue weight.

#### Catalase (CAT)

Cnidarians produce antioxidant enzymes as a mechanism to protect against reactive oxygen species generated by both hosts and pathogens during infection and associated immune responses.^78–80^ Catalase activity was measured using established protocols.^76,77^ Briefly, 5 µL of protein sample were combined with 45 uL of 50 mM PBS (7.0 pH) and 75 µL of 25 mM H_2_O_2_ in a UV 96-well plate. Immediately following addition of H_2_O_2_, absorbance at 240 nm was read every 45 seconds for 15 minutes. Catalase activity was determined from the steepest point of the reaction curve. Total hydrogen peroxide scavenged during this period was calculated using a standard curve of H_2_O_2_. Final catalase activity is reported as H_2_O_2_ scavenged per minute per mg of protein.

#### Total Phenoloxidase Activity (TPO)

The phenoloxidase pathway is an enzymatic cascade resulting in the synthesis of melanin – an effector molecule involved in wound healing and pathogen encapsulation.^78–80^ Here we quantify the phenoloxidase cascade using both total phenoloxidase activity (enzymatic activity of prophenoloxidase + phenoloxidase) and melanin concentration (see next section). Total phenoloxidase activity was estimated using modifications to existing protocols.^77^ Briefly, 20 µL of protein sample were combined with 20 µL of 50 mM PBS (7.0 pH) and 25 µL of 0.1 mg·mL^-1^ trypsin. Samples were then incubated on ice for 30 minutes to allow for cleavage of prophenoloxidase (PPO) into phenoloxidase (PO). We then measured total potential phenoloxidase activity (i.e. activity of any existing phenoloxidase in the sample + activity of prophenoloxidase activated during trypsin incubation). Following incubation, 30 µL of 10mM L-DOPA were added to each well to initiate melanin synthesis. Absorbance at 490 nm was read every 10 minutes for 4 hours. Total phenoloxidase activity was calculated from the steepest portion of the reaction curve and standardized to sample protein concentration.

#### Melanin (MEL)

Melanin tissue concentration was estimated using reserved tissue samples following established protocols.^76,77^ Briefly, samples were desiccated in a speed vac (Eppendorf Concentrator Plus) for a minimum of 12 hours. Following desiccation, samples were weighed to obtain total dry tissue weight. Then, 1.0 mm glass beads (Sigma Aldrich) and 400 µL of 10 M NaOH were added to each sample. Samples were vortexed for 20 seconds and stored in the dark for 48 hours. During this incubation, samples were vortexed briefly at 36 and 48 hours. At the end of this period, samples were centrifuged at 183 x *g* for 10 minutes at room temperature. A melanin standard processed identically to samples, was used to create a standard curve. To measure melanin concentration, 40 µL of the resulting supernatant from samples or melanin standard dilutions were transferred to half volume UV 96-well plates. Absorbance was measured at 490 nm and converted to melanin concentration using the standard curve. All melanin values were standardized by desiccated tissue weight.

#### Antibacterial Activity (AB)

Finally, we quantified antibacterial activity as a metric of the production of antimicrobial peptides and metabolites that can neutralize a wide variety of pathogenic bacteria.^78–80^ Antibacterial activity here was quantified by measuring bacterial growth rates of *Vibrio coralliilyticus* in the presence of protein extracts generated from host tissue. Antibacterial activity against *Vibrio coralliilyticus* BAA 450 was calculated using established protocols.^76^ Prior to the assay, samples were diluted to a standardized protein concentration. To initiate the reaction, 60 µL of sample (or buffer control) were combined with 140 µL of *V. coralliilyticus* diluted to an OD600 of 0.2. Absorbance was read at 600 nm every 10 minutes for 6 hours to generate bacterial growth curves. Percent inhibition of bacterial growth was calculated by comparing sample growth rate to growth rate of buffer controls from the steepest part of the curve.

### Energetic Assays

Total lipids and total carbohydrates assays were used to determine approximate host energetic budget. Lipids and carbohydrates are commonly used proxies for energetic budgets in multiple cnidarian species^81^. Total lipid and total carbohydrate assays were run in triplicate and duplicate, respectively, and measured in separate 96-well plates using the Cytation1 Cell Imaging Multimode Reader and Gen5 Software (Biotek). Tris + DTT buffer was used as a negative control for both assays.

#### Ash Free Dry Weight

Sample ash-free dry weight was calculated to standardize both lipid and carbohydrate assays using modifications from previous methods.^82^ All samples were weighed using an analytical scale. Briefly, a recorded volume of sample was added to tared aluminum boats and dried at 60°C in a drying oven for 48 hours. Samples were removed from the drying oven and weighed before combusting at 500°C for 4 hours in a furnace box. Following combustion, samples were weighed one final time. Ash-free dry weight was calculated as the average difference between the dry weights and ash weights of each sample and standardized to 125 µL volume.

#### Lipids

Lipid concentrations were estimated using a total lipid assay following established protocols.^83^ Samples were dried for 5 hours at room temperature in a speed vac and then combined with 500 µL 2:1 chloroform: methanol solution and 100µL of 0.05 M NaCl. Tubes were vortexed for 20 seconds before 2-hour incubation on a shaking incubator at 200 RPM, with brief vortexing approximately every 30 minutes. At the end of this period, samples were centrifuged at 1650 x *g* for 10 minutes at room temperature. One hundred microliters from the bottom organic layer of each sample were transferred in triplicate to a 96-well PCR plate. A serial dilution of corn oil standard dissolved in chloroform was also plated for the creation of a standard curve. Next, 50 µL of methanol were added to each well. The plate was incubated in a 70°C water bath for at least 15 minutes to evaporate the solvent, after which 150 µL of 18 M sulfuric acid (H_2_SO_4_) was added to each well. The plate was then covered and incubated at 90°C for 20 minutes and cooled to 4°C. Following this incubation, the plate was vortexed and 75 µL of each sample and standard were transferred into a new 96-well microplate. Following baseline absorbance measurement at 540 nm, 34.5 µL of mg·mL^-1^ vanillin in 17% phosphoric acid was then added to each well. After a 5-minute dark incubation period, absorbance values were read again at 540 nm and subtracted from the initial reads. Average differences in absorbance values were converted to lipid concentration using the standard curve and standardized by initial sample volume and ash-free dry weight.

#### Carbohydrates

Carbohydrate concentrations were estimated following established protocols.^84^ Fifty microliters of either sample or glucose standard were plated in duplicate on a 96-well microplate. Next, 150 µL of 18 M H_2_SO_4_ was added to each well, followed immediately by 30 µL of 5% phenol. Plates were incubated (uncovered) in a hot water bath for 5 minutes at 80°C and allowed to cool for 15-20 minutes. Absorbance values were measured at 485 nm and 750 nm. Total carbohydrates were calculated by converting the average difference between 750 nm and 485 nm absorbance values to glucose concentrations using a standard curve. Carbohydrate values were standardized to tissue weight using ash-free dry weight.

### Statistical Analyses

All statistical analyses were performed using R Version 4.4.0.^85^ Prior to hypothesis testing and modeling sample outliers were identified using the Rosner test from the EnvStats package (v2.8.1^86^) and removed. Shapiro-Wilke and Bartlett tests were used to confirm the assumptions of normal distribution and equal variances, respectively. Some assays failed to meet assumptions necessary for parametric testing, even following transformation, necessitating the use of non-parametric analyses.

#### Mortality

Survivorship data from our first subset of anemones were used to assess the effects of pathogen exposure, heat stress and clonal line on mortality. First, we tested for a significant effect of pathogen exposure on mortality using a Kaplan-Meier analysis (Survival ∼ pathogen challenge; n = 184; survival package v3.5.5^87^). Mortality data within pathogen exposed groups was then statistically evaluated using standard approaches for survival analyses, specifically a Cox proportional hazard’s model, which both allows for censorship of individuals (i.e. those that did not die) and incorporates data regarding rate of mortality.^88,89^

We investigated the interactive effects of heat stress and clonal line on mortality within the subset of pathogen-exposed anemones only (Survival ∼ temperature * clonal line; n = 92).

#### Univariate hypothesis testing

A second subset of anemones reserved for assessment of symbiont density, energetic reserves, and immune activity, was used to investigate hypotheses related to mechanisms of differential mortality. Given that there were no interactions between temperature treatment and clonal line in our cox proportional hazards modeling, we did not test for interactions in these physiological metrics. To isolate the effects of temperature from the effects of pathogen challenge, we separated out placebo samples and tested for differences in symbiont density (mean chlorophyll fluorescence intensity) or energetic reserves (total lipids and total carbohydrates) between temperature treatments (ambient vs. heat) and clonal lines within the placebo groups. The impacts of heat stress or clonal line on symbiont density, lipids, and carbohydrates were assessed independently with separate Welch’s two-sample t-tests (symbiont density and lipids) or non-parametric Wilcoxon Rank Sum tests (carbohydrates).

#### Redundancy analysis

To investigate the impact of our independent variables (temperature, pathogen exposure, and clonal line) and covariates (symbiont density, total lipids, and total carbohydrates) on host immune activity, we ran a redundancy analysis (RDA) using the vegan package (v2.6.6).^90^ Before starting with this multivariate approach, we tested for and confirmed no collinearity among our variables. Outliers in predictor variables were removed and all continuous variables were centered and scaled before separation into predictor (i.e., symbiont density, lipids, carbohydrates) and response (i.e., catalase, total phenoloxidase, and antibacterial activity, and melanin concentration) data frames. An RDA was used to identify which variables best predicted variation in immunity across our samples. We started by running an RDA with all possible predictor variables (i.e. clonal line, temperature treatment, pathogen exposure, symbiont density, lipids, and carbohydrates). From there, we used a forward selection to identify which predictor variables explained significant variation in our dataset. After identifying significant predictor variables, we ran a final RDA using the best-fit model determined by forward selection (∼ Treatment + Carbohydrates + Lipids).

#### Linear modeling

Finally, we used linear modeling to assess variation in individual immune parameters with respect to the statistically relevant factors determined via RDA (treatment, carbohydrate, lipids). For normally distributed immune parameters (catalase, total phenoloxidase, antibacterial activity) we used GLMs. Global models with all possible additive and interactive variables were analyzed for catalase, total phenoloxidase, and antibacterial activity independently. All models were dredged and ranked via AICc using the MuMIn package (v1.47.5)^91^, where applicable. Models with delta AICc values less than or equal to 2 were averaged to determine the best-fit models for each immune parameter. For melanin, which was right skewed with multiple zero measurements, we adopted a two-tiered strategy. First, we used a probit model (i.e. a generalized linear model with binomial error structure) to predict the presence of melanin among samples using heat treatment, carbohydrate and lipid concentration as predictors. We then removed samples in which melanin was not detected and used multiple regression to predict melanin concentrations employing the same predictors.

## Results

### Mortality

Anemone mortality was significantly greater in anemones exposed to the pathogen (Kaplan-Meier; *p* < 0.001; **Supplemental Fig. 1**). Of those anemones exposed to pathogen, 89% died during the 96-hour period, compared to 0% mortality within the placebo group. While there were no significant interactive effects of temperature treatment and clonal line on anemone mortality within pathogen exposed anemones controls (hazard ratio = 1.32; 95% CI [0.52 – 3.34]; *p* = 0.562), mortality rate was significantly greater in the heat stressed anemones relative to the ambient controls (hazard ratio = 2.29; 95% CI [1.11 – 4.75]; *p* = 0.0255) and among H2 anemones relative to VWB9 anemones (hazard ratio = 0.197; 95% CI [0.10-0.37]; *p* < 0.001; **Fig. 1; Supplemental Fig. 2**). When considering just those anemones exposed to pathogens, 82% of ambient and 97% of heat stressed individuals died within the 96-hour period. Furthermore, 100% of H2 and 82% of VWB9 anemones died.

**Figure 1:**
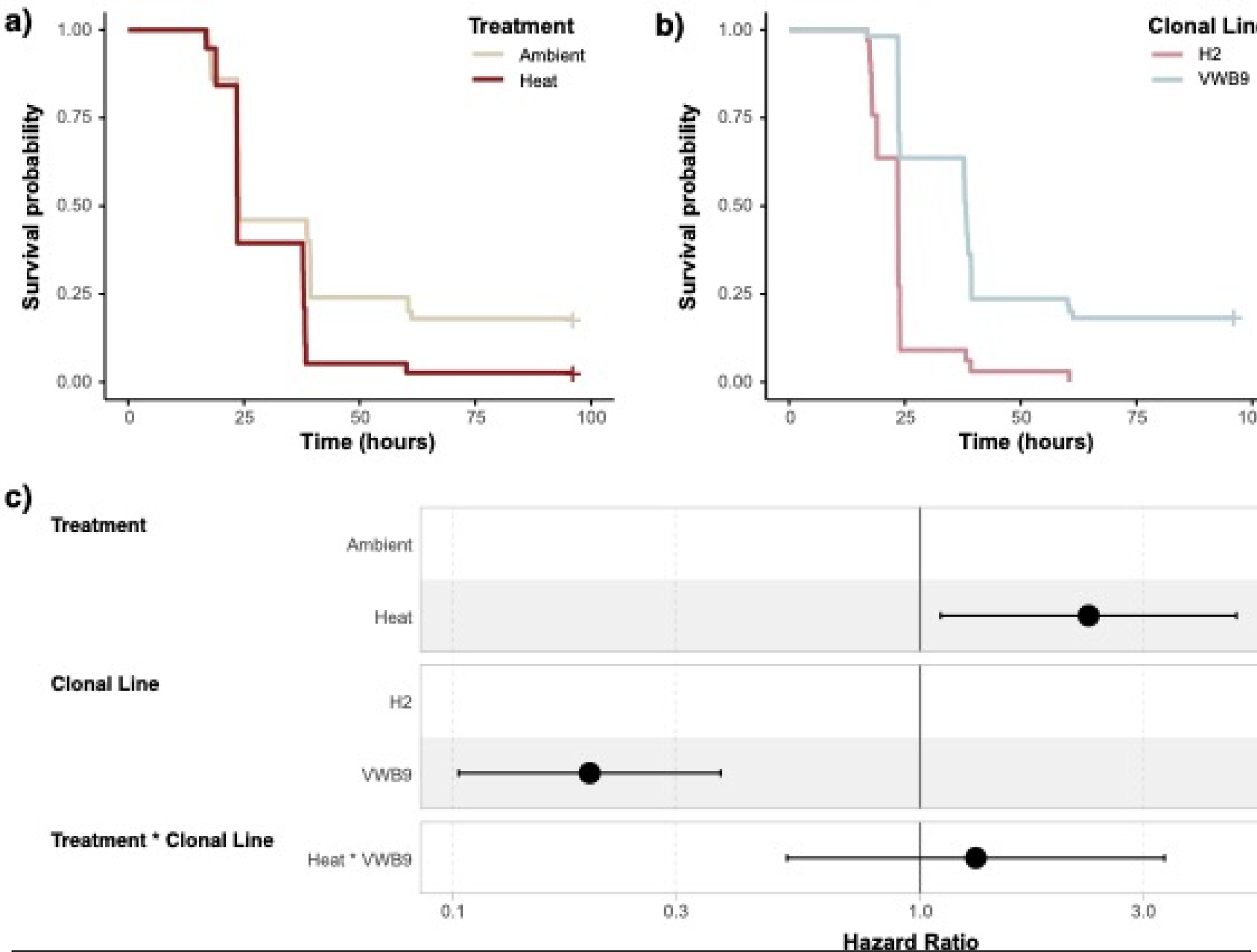
(a-b) Kaplan-Meier survivorship curves for pathogen exposed anemones split based on comparisons across (a) temperature treatment and (b) clonal line. (c) Cox proportional hazards models of pathogen exposed anemones compared across temperature treatment and clonal line. Vertical line at 1 indicates the reference to which the second group is compared. The distance from 1 indicates the probability of mortality (i.e. heat stress is about twice as likely to die from immune challenge as heat). Both treatment (p = 0.0255) and genotype (p < 0.001) had significant impacts on survival. Hazard ratios are displayed for both predictors and the interaction term with 95% confidence intervals.

### Symbiont Density & Energetic Reserves

Symbiont density, estimated by mean chlorophyll a fluorescence pixel intensity, was significantly higher in heat stressed anemones compared to ambient controls (Welch’s two sample t-test; t = −2.52, df = 40.181, *p* = 0.0157. 95% CI [-5617 – −620]; **Fig. 2a**). Symbiont density also varied across clonal lines (Welch’s two sample t test; t = 2.65, df = 38.3, *p* = 0.0118; 95% CI [768 – 5766]); H2 had significantly greater symbiont density than VWB9 anemones **(Fig. 2a**). For energetic reserves, total carbohydrates varied between treatment groups. Carbohydrates were significantly lower in heat treated anemones compared to anemones that remained at ambient temperature (Wilcoxon Rank Sum test; W = 392, *p* < 0.01; **Fig. 2b**), but not significantly different between clonal lines (Wilcoxon Rank Sum test; W = 220, p = 0.358; **Fig. 2b**). Total lipids did not significantly differ between either temperature treatment (Welch’s two sample t-test; t = −0.444, df = 40.12, *p* = 0.659, 95% CI [-0.143 – 0.0917], **Fig 2c**) or clonal lines (Welch’s two sample t-test; t = −1.27, df = 40.9, *p* = 0.213, 95% CI [-0.185 – 0.0424], **Fig 2c**).

**Figure 2:**
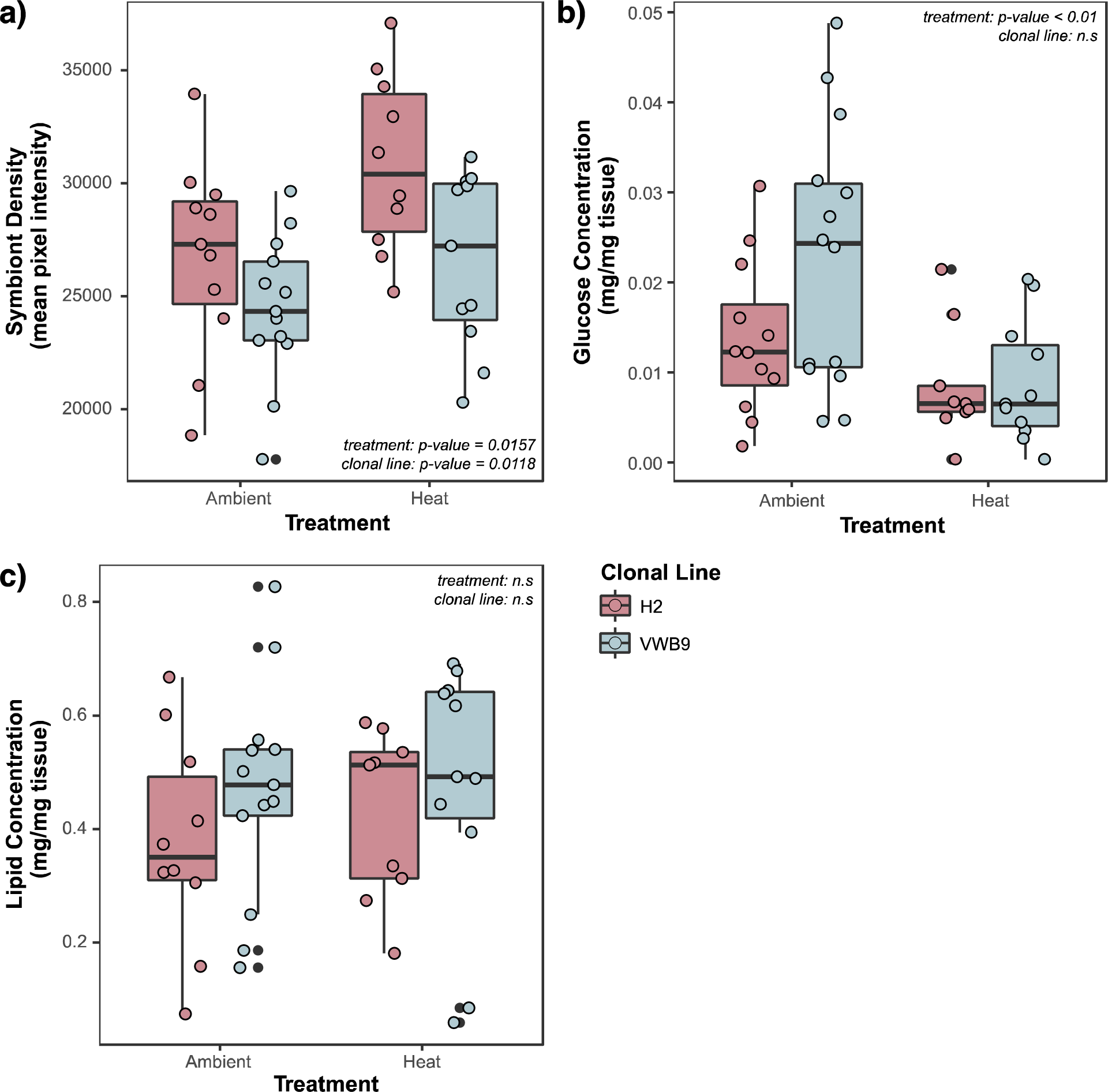
Box plots displaying **a)** symbiont density, **b)** lipid concentration, and **c)** carbohydrate concentration by temperature treatment and clonal line. Raw data values represented by overlaid points; boxes and points are colored based on clonal line. Significance (*p values*) for treatment and clonal line is reported.

### Immune Responses

The RDA explained significant variation along the first axis of the ordination, RDA1, which explained 18.12% of the total variation **(Fig. 3)**. Temperature treatment (ANOVA, *p* = 0.003), lipid concentration (ANOVA, *p* = 0.003), and carbohydrate concentration (ANOVA, *p* = 0.034), all explained significant variation in immune responses. Temperature treatment explained significant variation along both the RDA1 and RDA2 axes (**Fig. 3** boxplots; t-test, p < 0.0001). However, carbohydrate concentration explained variation mostly along the first RDA axis (**Fig. 3** regression bottom-left; Pearson correlation, p < 0.0001), while lipid concentration explained variation primarily on the second axis (**Fig. 3** regression top-right; Pearson correlation, p < 0.0001). The immune variables used to construct the ordination show strong association with the temperature treatments with all arrows representing the immune variables pointing towards the group of samples from the ambient treatment.

**Figure 3.**
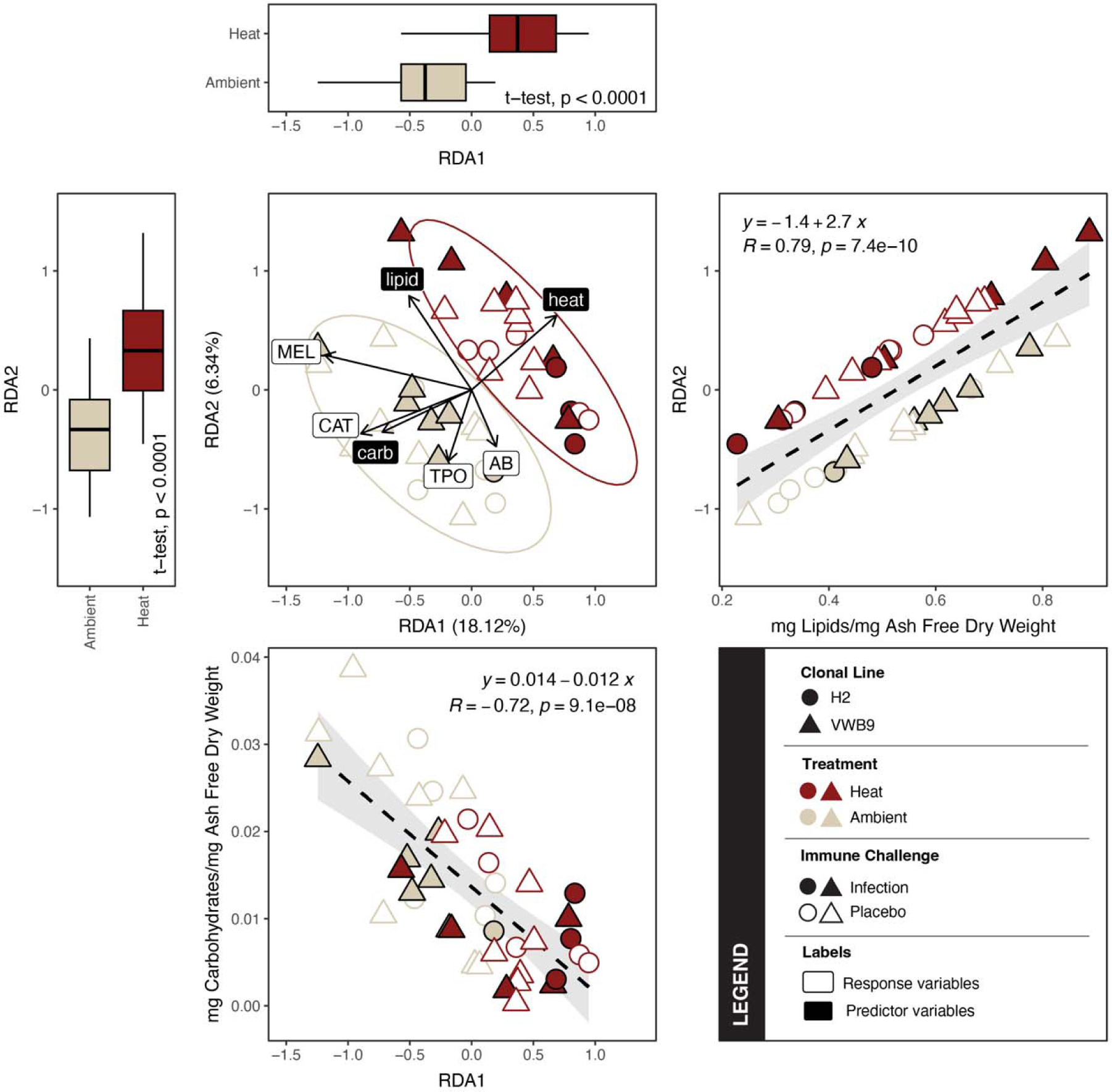
RDA plot (center) showing significant predictors (black labels) for explaining variation in immunity. Data points represent individual samples, and ordination was constructed using distances generated from immune parameters. Color indicates temperature treatment, shape indicates clonal line, and open shape/filled shape indicates immune challenge. Black labels represent predictor variables and white labels represent response variables. Arrows represent strength and directionality of each variable in multi-variate space. Boxplots on the top and far left indicate the distribution of heat and ambient points along the first and second axes to demonstrate significant variation explained by temperature treatment in each direction. Regressions on the bottom and far right show the correlation between carbohydrates and lipids on the first and second axes respectively to demonstrate significant explanation of variation by those variables on each axis. MEL: melanin; CAT: catalase; TPO: total phenoloxidase; AB: antibacterial activity.

Linear modeling of the effects of each of our significant predictors on individual immune metrics revealed metric-specific effects (**Table 3**). Three metrics of immunity (CAT, TPO, and MEL) were significantly lower in heat treated anemones relative to ambient temperatures (CAT GLM, *p* = 0.0129; TPO GLM, p = 0.0212; MEL zero-included binomial, *p* < 0.0430; **Fig. 4).** Additionally multiple metrics of immunity were positively associated with energetic measurements. Both CAT and MEL were positively associated with carbohydrate concentration (CAT GLM, *p* < 0.001; MEL zero-excluded GLM, *p* < 0.001; **Fig. 5**), and MEL was also positively associated with lipid concentration (zero-excluded GLM, *p* < 0.001; **Fig. 5**).

**Figure 4:**
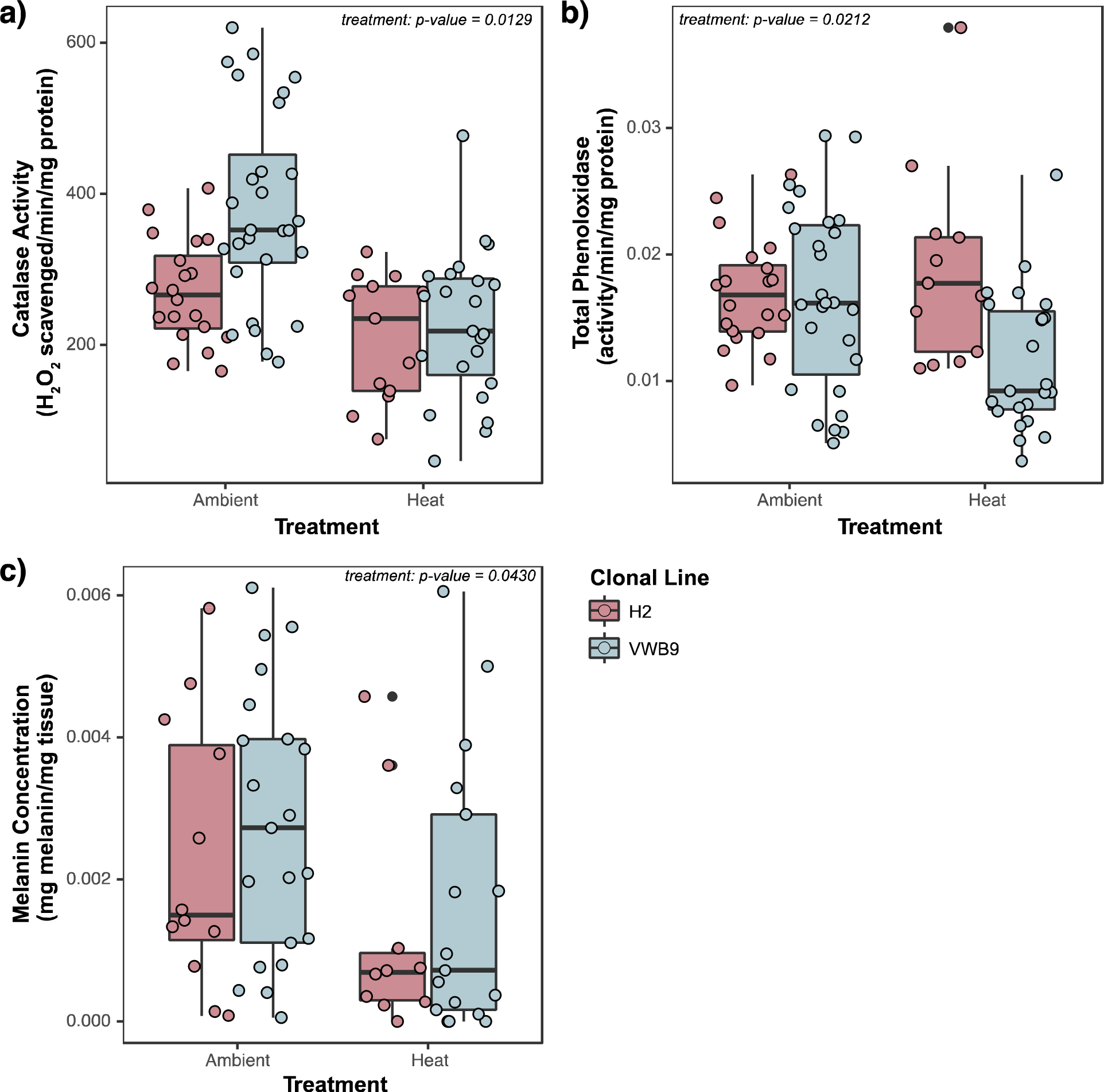
Box plot displaying differences in **a)** catalase activity, **b)** total phenoloxidase activity, and **c)** melanin concentration as a result of temperature treatment, divided by clonal line. Raw data values represented by overlaid points; boxes and points are colored based on clonal line. Significance (*p values*) for treatment is reported.

**Figure 5:**
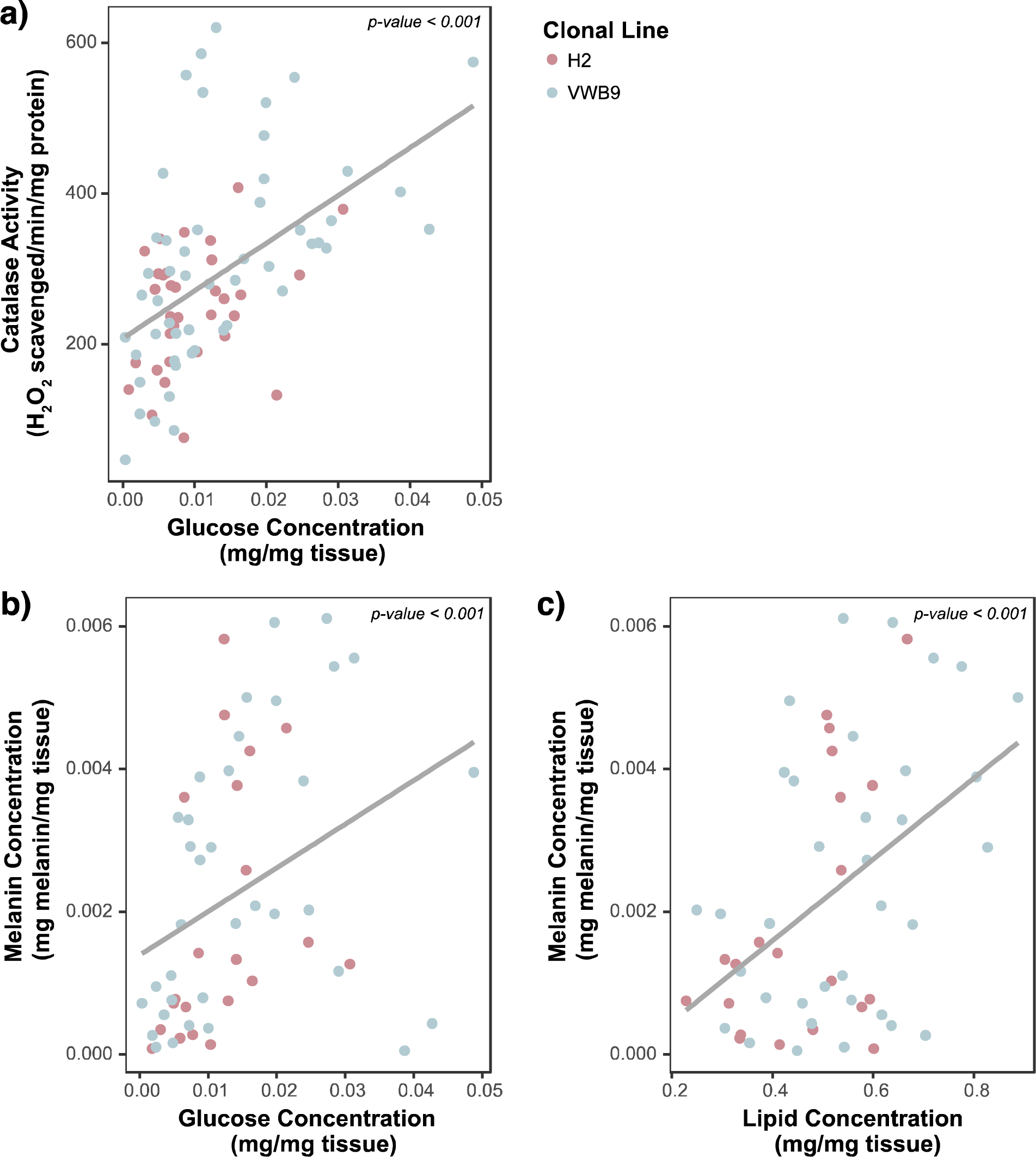
Best-fit linear models of the association between individual immune assays and energetic metrics: **a)** catalase activity and carbohydrate concentration, **b)** melanin concentration and carbohydrate concentration, and **c)** melanin concentration and lipid concentration Points represent individual samples color coded by clonal line. Trendlines represent linear models of association between immune and energetic metrics (independent of clonal lines). Significance (*p values*) for associations are reported.

**Table 3:**
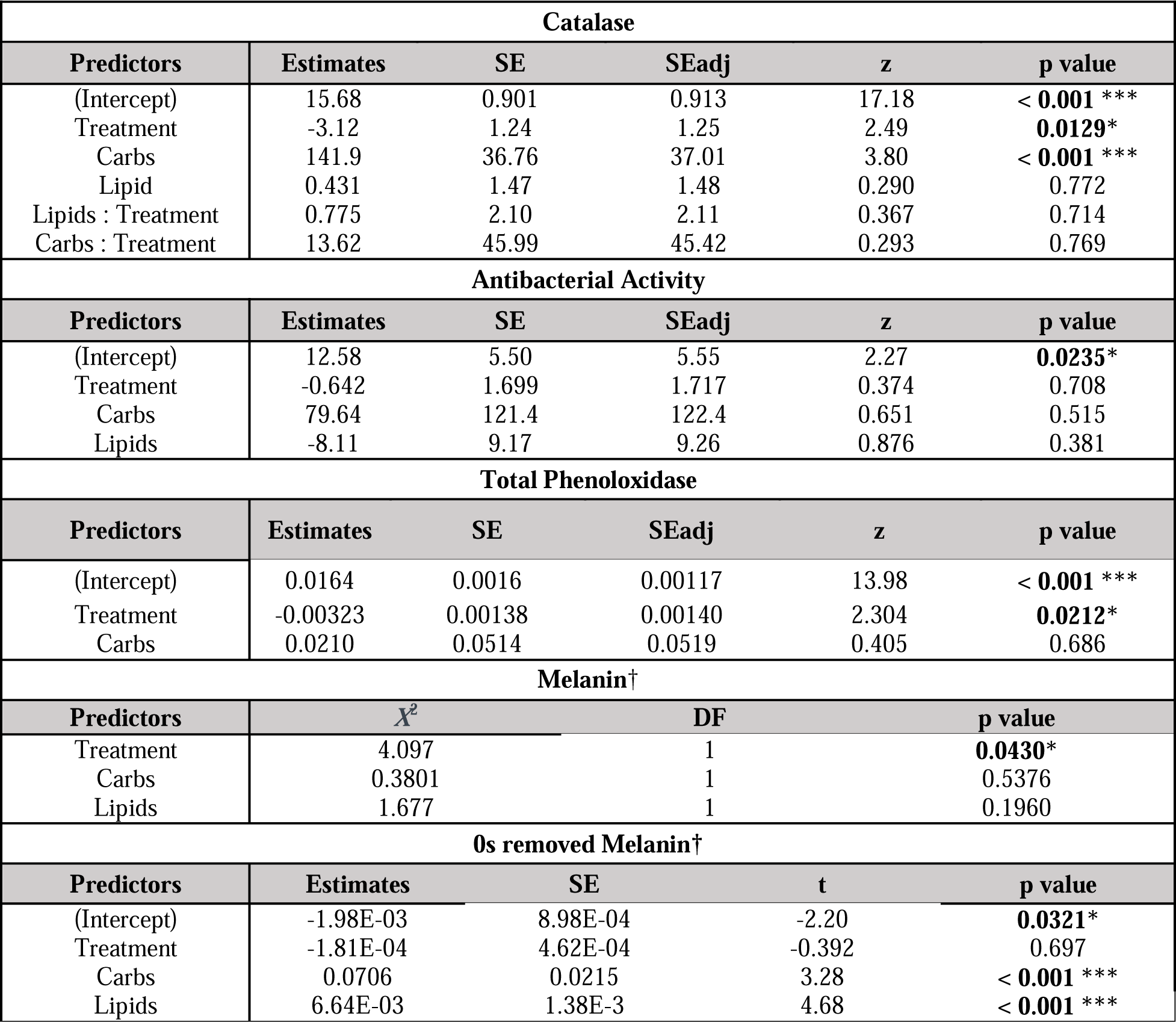
Best-fit linear models for each immune parameter when including temperature treatment, clonal line, immune challenge, and lipids as predictors. † indicate models that did not undergo model dredging and averaging. Asterisks (*) represent significant p-value (α = 0.05).

## Discussion

As anthropogenic climate change worsens, marine ecosystems are faced with a multitude of stressors that may form complex associations with each other.^3,5,17^ However, empirical study of interactive stressors, particularly those occurring asynchronously, has been limited, leaving many questions regarding the nature and mechanisms of associations amongst stressors.^92^ Cnidarians, which face a diversity of climate related stressors^21,80,93,94^, provide a relevant model to investigate the effects of multiple stressors (both interactive and sequential) on marine species. Here we conducted one of the first experimental studies investigating the consequences of prior heat stress on subsequent pathogen susceptibility. Our results support a long-held theory^95^ that previous heat stress increases later pathogen susceptibility and provide important new insight regarding these patterns. Specifically, we highlight changes in host cnidarian immunity following heat stress which may be driven by shifting resource allocation and likely contribute to observed patterns of differential pathogen susceptibility. These results are an essential first step towards understanding the cellular mechanisms which contribute to associations between sequential stressors.

### Mortality differs across temperature treatments and clonal lines

Susceptibility to pathogen exposure was significantly higher among anemones exposed to heat stress. This result is in agreement with documented increases in disease and disease-related mortality following coral bleaching in the wild.^24,40,44,96^ Notably, while previous laboratory experiments have documented impacts of simultaneous heat stress and pathogen exposure^45,47,50,51,97^ ours is the first to document the prolonged consequences of heat stress on pathogen susceptibility, more accurately representing observed environmental patterns in nature. Future studies combining our approach with emergent experimental disease techniques^98,99^ will allow for more broadscale applications to complex cnidarian and marine disease systems.

We also observed significant genotypic differences in mortality. Anemones from the VWB9 clonal line were less susceptible to pathogen-induced mortality than H2. Both clonal lines originate from similar geographic regions and host the same species of algal endosymbiont (*Brevolium minutum*)^100^, suggesting some degree of fine scale genetic, epigenetic, or plastic variation contribute to observed physiological differences between the lines. While intraspecific variation in cnidarian disease resistance has been linked to genetic variation in the wild^101,102^, there are a lack of studies investigating this variation in laboratory settings.^103^ Similarly, most experimental studies investigating pathogen dynamics in Aiptasia use only a single clonal line.^45,104–106^ Our findings suggest that inclusion of genetic variation in cnidarian studies, including those involving *E. diaphana*, may provide increased insight regarding the roles of genetic variability in contributing to stress resistance or susceptibility.

### Mechanisms of differential mortality: roles of symbiont density and energetic reserves

Beyond documenting differences in mortality as a result of environmental and genetic factors, we employed an integrative approach to investigate the underlying mechanisms driving these differences. Given the well-known contributions of symbiont density and energetic reserves to cnidarian host immunity and pathogen susceptibility^65,107,108^, as well as the documented effects of heat stress on these metrics^57,109–113^, we sought to examine differences in symbiont density and energetic reserves between temperature treatments. Contrary to expected results, symbiont density was higher in previously heat stressed individuals following two weeks of recovery, potentially due to *E. diaphana*’s capacity for exceptionally rapid recovery from dysbiosis.^75^ Thus, the two-week recovery between heat stress and pathogen challenge was likely ample time for anemones to recover from any dysbiosis and even overcompensate. Elevated symbiont densities in previously heat stressed individuals may be the result of unregulated symbiont growth during recovery. As negative associations have frequently been observed between symbiont density and immunity in other cnidarians^64–66^, the unchecked growth of symbionts during recovery may have contributed to differential mortality. Consistent with this hypothesis, H2 anemones, which were more susceptible to pathogens, also had consistently higher symbiont densities. In contrast to symbiont density, carbohydrate concentration was reduced in those anemones which had undergone previous heat stress. Thus, while symbiont density appears to recover quickly, carbohydrate concentration appears to recover more slowly, potentially contributing to increased pathogen susceptibility in previously heat stressed individuals. Time series and finer-scale approaches leveraging metabolomic or carbon-flux modeling are needed to determine the prolonged effects of heat stress on host energetic budget, and subsequent effects on host pathogen susceptibility.

### Mechanisms of differential mortality: roles of host immunity

Next, we focused on the potential roles of shifts in immunological activity which might contribute to observed patterns of mortality. When considering immunity using multivariate statistics, we saw significant effects of temperature, carbohydrate and lipid concentration on immune response. Particularly, we saw indication of immune suppression in anemones that had undergone prior heat stress. Examination of the effects of heat stress on individual immune parameters revealed strong suppression of catalase activity, total phenoloxidase activity, and melanin concentration. While numerous studies have evidenced the short term effects of heat stress (days – weeks) on cnidarian host immunity^48,114–116^ the long-term effects of heat stress (months – years) on immune activity are not well documented.^117,118^ Many studies have noted heat-induced activation of a variety of immune parameters, including antioxidant (e.g., catalase) and phenoloxidase enzymes^48,114,117^, though some studies report minimal or negative impacts of heat on immunity.^52–54^ Positive associations between heat stress and immune responses during simultaneous stressors may be driven by overlap between response mechanisms; some heat stress biomarkers (ie HSP70) are known to activate immune pathways such as prophenoloxidase.^119,120^ In contrast to the synergy which often occurs between heat stress and immune activation during simultaneous stressors, we document pronounced suppression of immunity (catalase, phenoloxidase, melanin) after two weeks of recovery from heat stress. Our approach suggests significant immune suppression occurs during, or as a result of, recovery from heat stress, though the exact timing is unclear. Frequent, repeated sampling during heat stress and recovery will improve understanding of the prolonged effects of increasing SSTs on cnidarian immunity.

### Investigating the role of symbiotic immunosuppression

While immune suppression following heat stress may be the result a number of mechanisms, one of the most commonly proposed mechanisms involves dynamic changes in symbiosis. While our observation of higher symbiont density in groups more susceptible to pathogen challenge (i.e. previously heat stressed and H2 anemones) is consistent with this hypothesis, we did not document any significant associations between symbiont density and our measured metrics of immunity. Two potential explanations exist: 1) we measured a limited suite of immune metrics; symbiont-induced immune suppression may have effects on unmeasured metrics, or 2) indirect or time lagged effects of changing symbiont densities may be influencing observed patterns of differential mortality. Specifically, rapid and excessive symbiont proliferation (evidenced by higher symbiont densities in heat treated anemones) may have had significant prolonged effects on host immunity that are not detected at the time of sampling (i.e. time lagged effects). Future studies incorporating more immune parameters and time-series approaches to sampling may be used to test these hypotheses.

### Investigating the role of energetic immunosuppression and the importance of resource allocation

A second potential hypothesis to explain observed associations between heat stress and immunity is that changes in resource allocation as a result of resource limitations during dysbiosis may result in immune suppression. The strongest support for this hypothesis comes from our analyses involving carbohydrate concentrations, which were significantly reduced as a result of prior heat stress. Additionally, carbohydrate concentration was strongly positively associated with both catalase activity and melanin concentration, both of which were more active/abundant in samples from the ambient group. Combined, these data suggest that reductions in carbohydrate concentrations due to prior heat stress have direct effects on immunity; as energetic reserves decrease in response to prior heat-stress certain immune responses appear suppressed. These findings are consistent with synthesis of previous findings. Heat stress both causes reallocation of host energetic reserves to stress responses, and often impacts overall energetic reserves through the reduction in symbiont density which provide nutritional resources to hosts.^113,121–126^ This loss of energetic resources likely results in a decrease in the total energy available to allocate to immune defenses, consistent with resource allocation theory.^61,62,127^

Associations between lipid and melanin concentrations further support the notion that changes in energy affect host immunity. Melanin concentration was also significantly positively correlated to lipid concentration, which was not affected by prior heat stress. Reduced effects of prior heat stress on lipid compared to carbohydrate are consistent with previous findings; stored carbohydrates, rather than lipids, are preferentially used as an energy source by hosts during bleaching recovery in other cnidarians.^113^ The association between melanin and lipids, which were relatively unaffected by prior heat stress, suggests that melanin production potentially draws resources from both lipids and carbohydrate sources. The use of both lipids and carbohydrates to fuel melanin production could account for comparatively reduced impacts of heat on this metric compared to other immune parameters like catalase. Together, our results suggest that changes in total energetic resources directly impact host immunity in the aftermath of heat stress.

### Conclusion & Future Directions

Multiple stressors are a common consequence of anthropogenic climate change and can have pervasive effects on a variety of marine ecosystems.^17^ As such, an improved understanding of organismal responses to climate-induced stressors is imperative to the conservation of these ecosystems. Here we detail one of the first studies to link prior heat stress and increased disease susceptibility in a marine species. Our study establishes the ability to reproduce heat stress and disease associations under laboratory conditions using a tractable model system, allowing for more nuanced and robust investigations of ecologically relevant phenomena. Using this experimental approach, we were able to demonstrate prolonged effects of prior heat stress on host immunity, which may contribute to observed increased pathogen-induced mortality following recovery from heat stress. Furthermore, our results implicate changes in resource availability as a potential mechanism of heat-induced immunosuppression. Further studies incorporating time series approaches will be necessary to fully disentangle the observed patterns and associated mechanisms and assess their broad applicability to marine systems. Specifically, future work should aim to sample across multiple timepoints from the onset of bleaching through acute and long-term recovery to determine the effects of dynamic changes in symbiont density and energetic reserves on host immunity and pathogen susceptibility. This, coupled with more nuanced approaches such as fine-scale metabolomics and transcriptomics, should provide a clearer picture of the nuances of changing resource allocation, symbiont density, and immunity which link prior heat stress to increased pathogen susceptibility in corals. In summary, our results provide an important starting point for understanding the links between heat stress and disease that will aid in the conservation of marine species, including cnidarians, in the face of a changing climate. Most importantly, they generate testable hypotheses that will be invaluable in better understanding the effects of multiple stressors on diverse species and ecosystem functions.

## Data and Code Availability

All raw data and R code used in described analyses is supplied as supplementary files.

## Conflict of Interest

The authors declare they have no competing interests.

## Notes

### Competing Interest Statement

The authors have declared no competing interest.

### Summary of Updates

Response to reviewers and updated statistical analyses

