## Supplementary figures and images for "Prior heat stress increases pathogen susceptibility in the model cnidarian *Exaptasia diaphana*"

### Supplemental Fig. 1

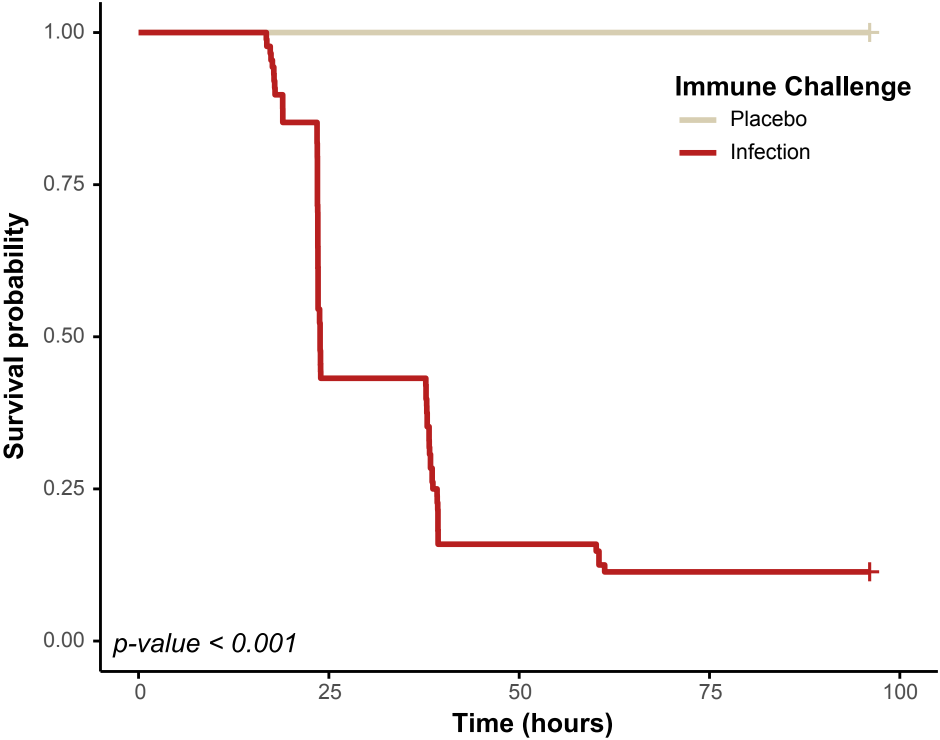

### Supplemental Fig. 2

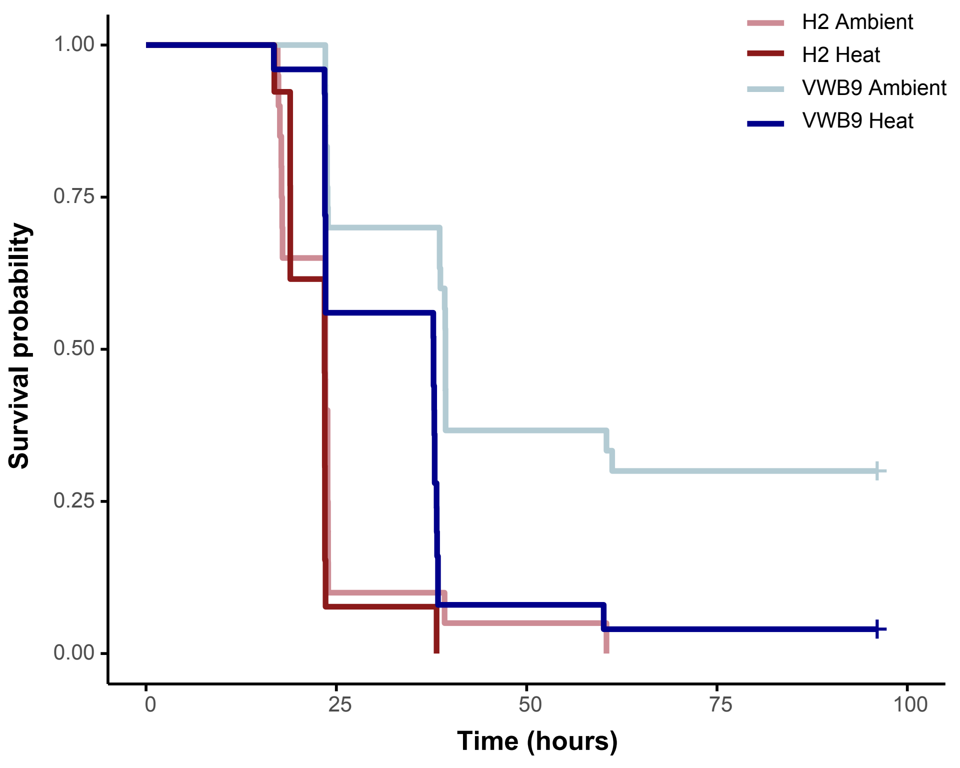
